# Optimization and Comparative Evaluation of SimpliAmp and Bio-Rad Thermal Cyclers for PCR-Based Detection of *Vibrio cholerae*

**DOI:** 10.1101/2025.10.17.683026

**Authors:** Victor Okpanachi, Khai Truong, Gerardo Lopez, Kerry K. Cooper, Aminata Kilungo

## Abstract

Cholera continues to pose a significant public health risk in low- and middle-income countries (LMICs), necessitating quick, cost-effective, and dependable diagnostics for effective outbreak surveillance and management. In this study, we optimized and tested two PCR-based assays targeting a uniplex assay with the *ompW* gene, and a multiplex assay with *ctxA, O1-rfb*, and *O139-rfb* genes of *Vibrio cholerae (V. cholerae)* on two thermal cycler platforms with different cost: the Bio-Rad C1000 Touch, which is more expensive, and the SimpliAmp, which is cheaper. Cycling conditions, annealing temperatures, and primer concentrations were systematically optimized to improve efficiency and reagent use. We tested our assay sensitivity by serially diluting gBlock DNA from 2000 to 50 copies. Quo data software was used to estimate the limit of detection and Fisher’s exact tests with Holm–Bonferroni correction to assess any differences (p<0.05) across all dilution levels on both platforms. The *ompW* assay achieved identical performance on both platforms, with 100% detection down to 250 copies and an LOD_₉₅_ of 523 copies (95% CI: 347– 786; p = 1.000). For the multiplex assay, *O1-rfb* showed slightly higher sensitivity on SimpliAmp (LOD_₉₅_= 509 copies, 95% CI: 339–766) than on Bio-Rad (LOD_₉₅_ = 333 copies, 95% CI: 255–688; p = 0.217). The *ctxA* assay had an LOD_₉₅_ of 684 copies (95% CI: 501–1323) on SimpliAmp and 974 copies (95% CI: 714–1784) on Bio-Rad (p = 0.089). For *O139-rfb*, SimpliAmp and Bio-Rad achieved LOD_₉₅_ values of 639 (95% CI: 490–1251) and 1060 copies (95% CI: 719–1562), respectively (p = 0.179). In general, the differences in performance between platforms were not statistically significant. These results highlight the potential of SimpliAmp as a reliable, low-cost alternative for cholera diagnostics in LMICs and low-resource settings, supporting broader access to molecular tools for outbreak investigation and public health surveillance.

## Introduction

Cholera is a serious diarrheal disease that causes dehydration quickly and is one of the fastest fatal infections if not treated right away (1). Its causative agent is a motile, facultative anaerobic, comma-shaped, Gram-negative bacterium, *Vibrio cholerae,* which naturally exists in aquatic environments (2). The O1 and O139 strains are primarily accountable for significant outbreaks and epidemics among its various strains. The cholera toxin encoded by the ctxA gene is a key virulence factor for these strains, playing key roles in disease pathogenesis (3). Together, these virulence strains contribute to the organism’s ability to persist in the environment and drive the recurrent global pandemics associated with *V. cholerae*.

Cholera continues to spread globally and has been responsible for seven pandemics (4). Based on the structure of the lipopolysaccharide, more than 200 serogroups of *V. cholerae* have been identified (1). Among these serogroups, O1 and O139 have been linked with cholera epidemics (3,5). Given that several serogroups of *V. cholerae* have been identified, it is the evolution and global spread of specific epidemic strains—particularly the O1 serogroup—that have shaped the trajectory of cholera pandemics, paving the way for their emergence and prevalence till the current seventh cholera pandemic is attributed to strains from the seventh pandemic El Tor (7PET) lineage.

At present, there are three primary methods for detecting *V. cholerae*: (1) Traditional culturing: which allows for the isolation of viable bacteria that can be grown for further testing. However, it has several drawbacks, including being time-consuming, labor-intensive and risk of contamination (6,7). (2) Time-of-flight mass spectrometry: which can analyze bacteria-specific proteins within a short time but, obtaining bacterial monoclonal strains involves isolating the culture(6), and (3) Molecular biology techniques such as polymerase chain reaction (PCR), which target specific nucleic acid sequences and facilitate more rapid, sensitive, and precise detection of V. cholerae. It also comes with drawbacks including higher cost compared to the traditional culture methods, tendency to produce false positive and negative results (6,8). PCR is especially useful for diagnosing infectious diseases like cholera due to its ability to amplify specific DNA segments associated with toxigenic traits (9,10). By targeting genes like *ctxA* and the *O1-rfb* and *O139-rfb* strains, PCR ensures faster identification of toxigenic *V. cholerae* from clinical and environmental samples.

PCR is a molecular technique that has revolutionized infectious disease diagnostics due to its ability to amplify specific DNA segments with remarkable sensitivity, enabling the detection of even trace of numerous pathogens including bacteria, viruses and fungi (9–13). Its strong diagnostic strength has made it a valuable tool in clinical microbiology, epidemiology, and environmental health. The rapid turnaround time and potential for automation in PCR processes also enhance clinical workflows, providing timely and accurate results crucial for patient management. Given that importance, PCR requires continuous improvement and adaptability to maintain reliability, accuracy, and accessibility—particularly in low-resource settings where its benefits are most needed. contamination leading to false positives, the need for careful primer design and reaction optimization, and potential non-specific amplification in multiplex assays are still some of the challenges that can be encountered (14–16). Despite the remarkable diagnostic power of PCR, its reliance on expensive laboratory equipment and reagents continues to limit accessibility and widespread use in low-income and resource-poor settings (15–18). The conventional thermocyclers, such as the Bio-Rad platform, are costly for many LMICs, which prevents routine testing in these settings where it is most needed. Even though several molecular assays have been developed to enhance the detection of infectious diseases, their implementation is frequently limited in LMICs by infrastructural and financial barriers to purchasing conventional thermocyclers such as the Bio-Rad. Addressing these barriers is necessary in ensuring that this cornerstone of molecular diagnostics continues to deliver accurate and equitable results across diverse settings.

Irrespective of the platform, robust amplification depends on standardized reaction protocols, which are critical for ensuring efficiency and reproducibility across laboratories (19). While several studies have demonstrated that Bio-Rad platforms provide a reliable framework for *V. cholerae* diagnostic (20,21), others have emphasized their high accuracy in quantifying nucleic acids. For example, Chen and others in 2019 noted the use of Bio-Rads technologies in detecting specific mutations circulating cell-free DNA, highlighting the platform precision in molecular diagnostics (22). This level of precision is beneficial in applications where genomic integrity is critical, making the Bio-Rad thermal cyclers like the C1000 Touch a solid choice for laboratories aiming for high compliance with diagnostic standards. However, despite its proven reliability, no study to our knowledge has directly compared the performance of such high-cost systems with low-cost alternatives for V. cholerae detection. This study addresses that critical gap by evaluating the comparability of the Bio-Rad C1000 Touch (**Fig.1**) with the more affordable SimpliAmp platform by Applied Biosystems® (**Fig.2**), which costs nearly 50% less. Demonstrating their performance will provide compelling evidence on whether the less expensive platform is a feasible and effective PCR solution for LMICs, where affordability remains a key barrier to infectious disease monitoring.

**Fig. 1:**
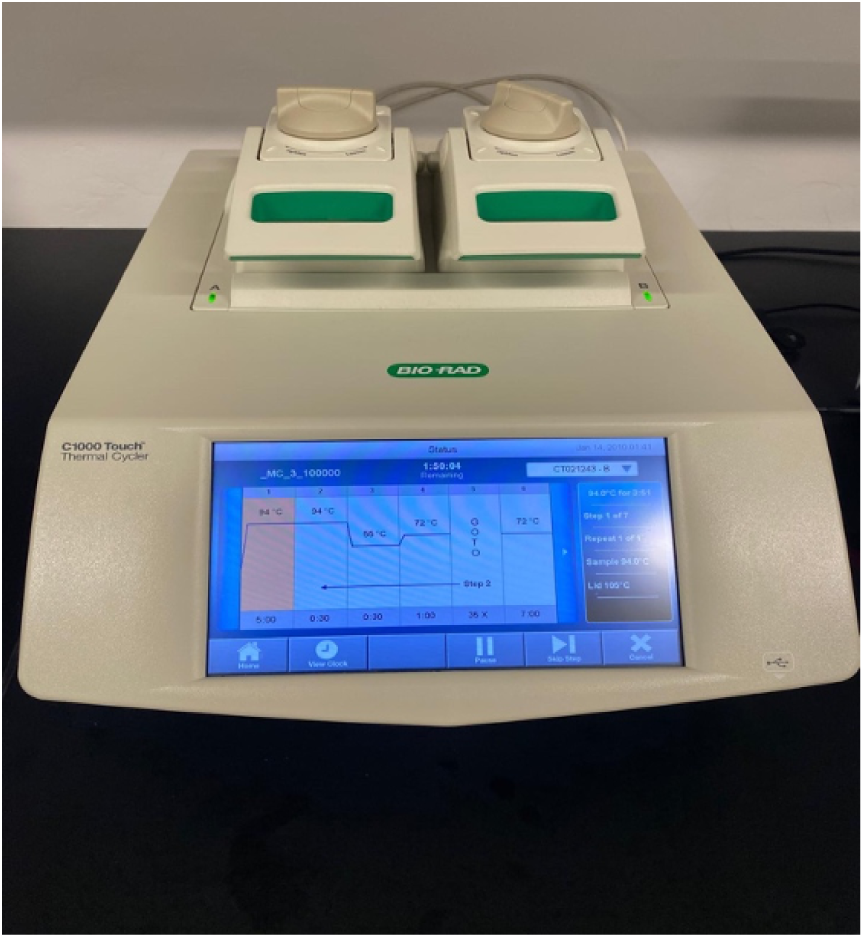
Bio-Rad C1000 Touch Thermal

**Fig. 2:**
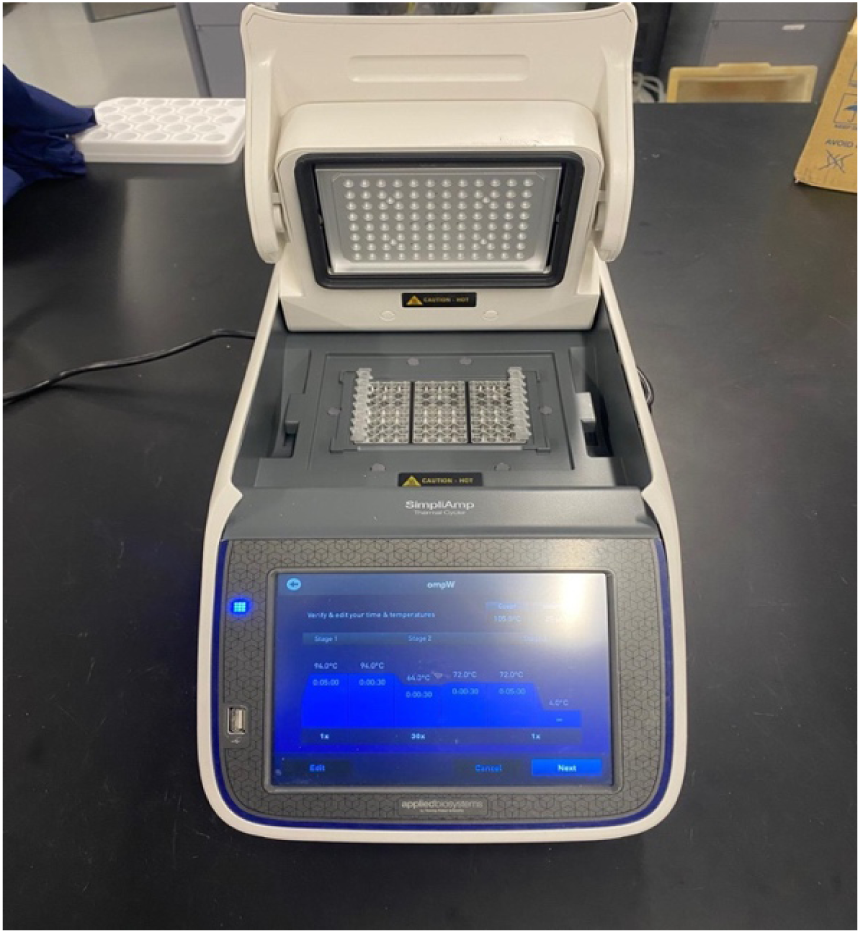
SimpliAmp Thermal cycler

Therefore, this study aims to optimize a PCR-based molecular assay for the detection of *V. cholerae* on both the Bio-Rad and SimpliAmp platforms. The specific objectives of this study are: (1) determine ideal cycling conditions (cycling time and annealing temperatures); (2) evaluate and determine an ideal primer concentration of the limit of detection (LOD) for the detection of singleplex *ompW* and multiplex *ctxA*, O1-*rfb*, and O139-*rfb* assays on both platforms; and (3) determine how effective both platforms are at detecting these targeted strains.

## Methods and procedure

### Selection of *V. cholerae* Target Genes

To detect the presence of *V. cholerae*, conventional polymerase chain reaction (PCR) was used in analyzing the presence of the *ompW* gene encoding the outer membrane protein in *V. cholerae* as described by Nandi and others, (2000) where they developed a rapid method for species-specific identification of *Vibrio cholerae* using primers targeted to the gene of outer membrane protein *OmpW* (23). We selected the *ompW* marker gene due to its demonstrated specificity for *V. cholerae*. This study established that the *ompW* gene is a reliable target for the accurate identification of *V. cholerae* strains, distinguishing them from other related species (23).

Three additional genes were targeted for amplification using PCR, including *ctxA,* the cholera toxin A subunit (*ctxA*), and two pandemic specific serogroups O1(O1-*rfb*) and O139(-O139-*rfb*) as described by Alam and others(24). These serogroups were selected because they are the only serogroups currently associated with cholera epidemics. Simultaneous amplification of these three primer pairs (forward and reverse) targeting the O1-*rfb*, O139-*rfb*, and *ctxA* genes was performed using synthetic double-stranded DNA fragments (gBlocks; Integrated DNA Technologies, USA) in 0.2 ml PCR tubes. The primer sequences and the expected amplicon length of the four genes are shown in (**Table 1**).

**Table 1:**
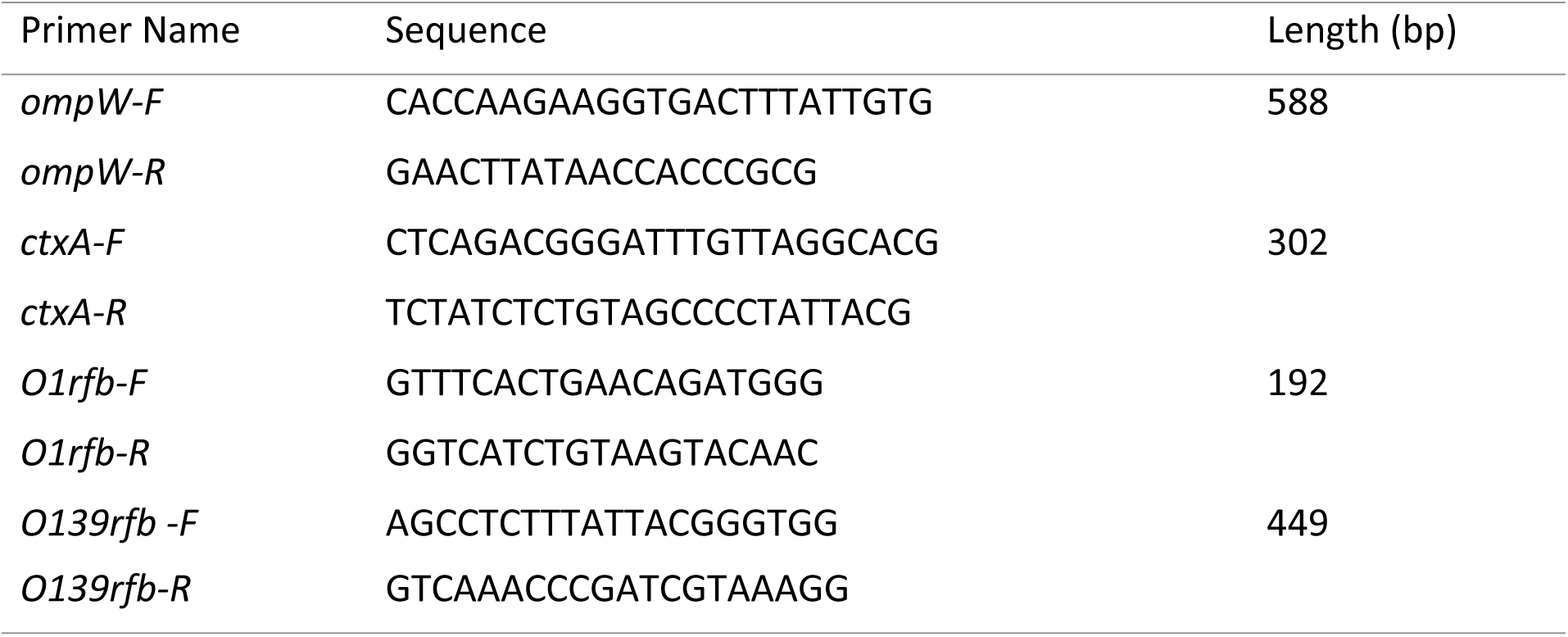
Summary of primer sequences and amplicon length

### Uniplex PCR assay of *ompW* gene

The uniplex PCR assay was conducted using gBlocks samples as the source template and nuclease-free water as the no template control. The positive control was spiked with 100,000 copies of the *ompW*_AA gBlock (IDT, USA), a synthetic ultramer representing the 588 bp *V. cholerae ompW* amplicon in which we had engineered a traceable double “AA” mutation to differentiate the synthetic DNA from natural *V. cholerae* DNA. PCR conditions used were previously described by Nand et al. and considered as baseline (23), indicating a 25 µL reaction containing 0.30 µL (5 U/µL) of Ex Taq DNA Polymerase (Takara Bio, USA), 2.50 µL of a 10X reaction buffer containing 20mM MgCl_2_ (Takara Bio, USA), 2.50 µL of 2.5mM dNTPs, 2.50 µL of 10 µM forward and reverse primers each, 11.80 µL of nuclease-free water and 2 µL of synthetic DNA template. The PCR cycling conditions began with an initial denaturation step at 94°C for 5 min, followed by 30 cycles of denaturation (94°C, 30 sec), annealing (64°C, 30 sec), and extension (72°C, 30 sec), and a final extension step at 72°C for 7 min as the baseline conditions.

### Multiplex PCR of *ctxA*, O1-*rfb*, and O139-*rfb*

A multiplex PCR assay was conducted using gBlocks as the source template and nuclease-free water as the no template control. The positive control was spiked with 100,000 copies of the *ctxA_AA*, *O1-rfb_AA*, and *O139-rfb_AA* gBlocks (IDT, USA), synthetic ultramer representing 302 bp, 192 bp, and 449 bp segments, respectively, each of which we had engineered with traceable double “AA” mutations to distinguish them from natural DNA. PCR conditions described by Alam et al in their study of toxigenic *V. cholerae* in aquatic environment (24), where used in this study and considered as baseline. Briefly, a 30 µL reaction consisted of 0.15 µL (5 U/µL) of Ex Taq DNA polymerase (Takara Bio, USA), 3.00 µL of 10X reaction buffer, 1.8 µL of 25mM MgCl_2_, 1.02 µL of 5 µM *ctxA* of each primer, 1.62 µL of 5 µM O1-*rfb* of each primer, 1.62 µL of 5 µM O139-*rfb* of each primer, 11.01 µL nuclease-free water and 2 µL of synthetic DNA template was used. The PCR cycling conditions began with an initial denaturation step at 94°C for 5 min, followed by 35 cycles of denaturation (94°C, 1 min), annealing (55°C, 1 min), and extension (72°C, 1 min), and a final extension step at 72°C for 7 min were used as the baseline conditions.

### Optimization of the uniplex *ompW* and multiplex *ctxA*, O1-*rfb*, and O139-*rfb* PCR assays

PCR optimizations for both singleplex *ompW* and the multiplex *ctxA*, O1-*rfb*, and O139-*rfb* were carried out simultaneously on the SimpliAmp thermocycler by Applied Biosystem® and Bio-Rad C1000 Touch thermal cycler by Bio-Rad Laboratories, Inc. Cycling time and annealing temperatures were tested for both assays. In addition, primer concentrations were also tested for both assays with the possible goal of saving reagent cost for LMICs. The summary of all optimization steps is displayed in **Fig 3** below.

**Fig. 3:**
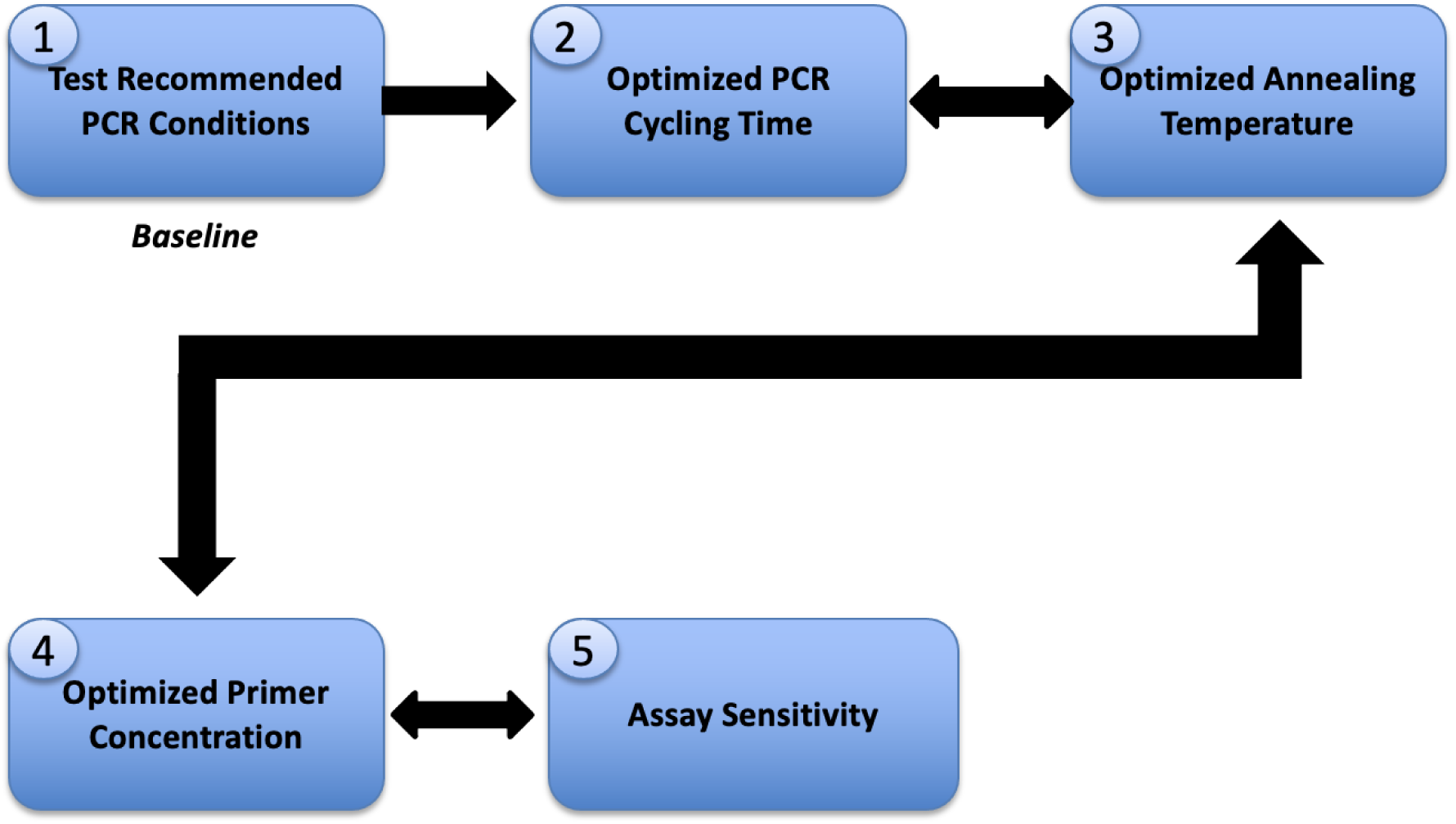
Methodology flowchart for molecular assay optimization. The baseline assay (1) was first tested under recommended PCR conditions. Optimization steps (2–4) were then performed sequentially, beginning with cycling times, annealing temperature, and primer concentration. Sensitivity evaluation (5) was conducted as the endpoint to compare baseline and optimized conditions.

For *ompW*, three different cycling conditions were tested. The first cycling condition included denaturation (94°C, 30 sec), annealing 64°C, 30 sec) and extension (72°C, 30 sec) that was from a baseline study (23), and the summary of remaining two optimized cycling conditions are summarized in (**Table 2**). Using gradient PCR on both thermal cyclers, annealing temperatures for *ompW* PCR assay were investigated from 61°C-68°C with a one-degree increment in a stepwise manner until a relatively strong and equal bands on both SimpliAmp thermal cycler and Bio-Rad C1000 Touch platforms were achieved. Additionally, we adjusted primer concentrations for *ompW*-F and *ompW*-R between 1.00 µM to 0.60 µM at 0.10 µM increment for both platforms until a clear visible band was obtained. We assessed each concentration combination after gel electrophoresis by visually examining if the target band was clear enough. The combination that yielded clear and equal bands for all the PCR targets, with the lowest primer concentration, was selected as our best overall combination given that one of our objectives is to reduce costs.

**Table 2:**
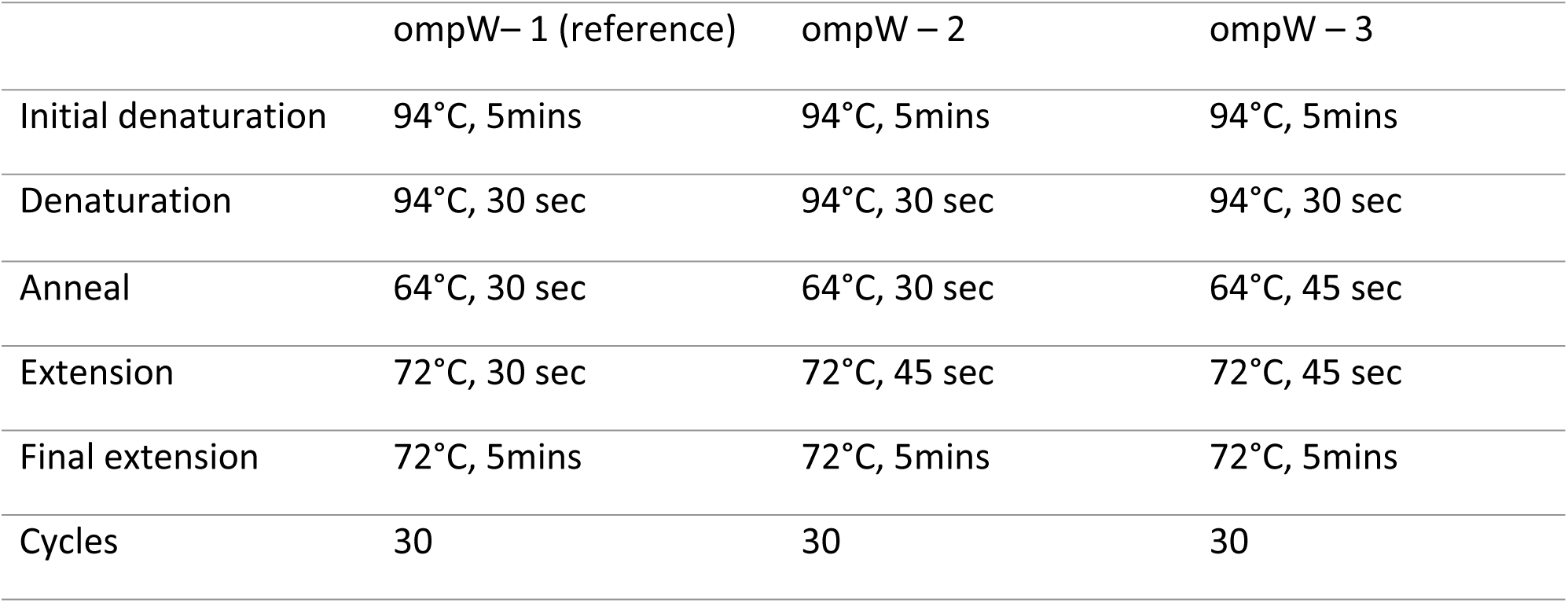
Summary of ompW assay cycling time optimization

For the multiplex cycling conditions, four cycling conditions were tested. The first PCR was from a baseline study (25), denaturation and annealing time were reduced to (94°C, 30 sec) and (55°C, 30 sec) respectively while, extension time was reduced to (72°C, 45 sec). Summary of all conditions, adjusted for different denaturation, annealing and extension cycling time are summarized in (**Table 3**). Like *ompW* assay, annealing temperatures for the multiplex PCR assay were adjusted from 55°C - 60°C with a one-degree increment in a stepwise manner until a relatively strong and equal bands on both SimpliAmp and Bio-Rad C1000 Touch platforms were achieved. For the multiplex assay, different primer concentration combinations for *ctxA* (0.17 µM, 0.15 µM and 0.13 µM) and both O1 and O139 (0.27 µM, 0.20 µM and 0.10 µM) were evaluated in a matrix format (**Table 4**) to determine the most suitable combination that can detect down to lower copies on both platforms.

**Table 3:** Summary of multiplex assay cycling time optimization

|  | Multiplex–1 (reference) | Multiplex – 2 | Multiplex– 3 | Multiplex– 4 |
| --- | --- | --- | --- | --- |
| Initial denaturation | 94°C, 5mins | 94°C, 5mins | 94°C, 5mins | 94°C, 5mins |
| Denaturation | 94°C, 1min | 94°C, 30 sec | 94°C, 30 sec | 94°C, 45sec |
| Anneal | 55°C, 1min | 55°C, 30 sec | 55°C, 30 sec | 55°C, 45 sec |
| Extension | 72°C, 1min | 72°C, 45 sec | 72°C, 1 min | 72°C, 45 sec |
| Final extension | 72°C, 7mins | 72°C, 7mins | 72°C, 7mins | 72°C, 7mins |
| Cycles | 35 | 35 | 35 | 35 |

**Table 4:** Matrix combinations for multiplex assay primer concentration (µM)

|  |  | *ctxA* Primers |  |  |  |  |
| --- | --- | --- | --- | --- | --- | --- |
|  |  | 0.17 | 0.15 | 0.13 |  |  |
| 01-rfb | 0.27 | 1 | 2 | 3 | 0.27 | 0139-rfb |
|  | 0.2 | 4 | 5 | 6 | 0.2 |  |
|  | 0.1 | 7 | 8 | 9 | 0.1 |  |

### Visualization of PCR products

After amplification, the PCR products were visualized using gel electrophoresis with a 1.7% agarose gel stained with Invitrogen SYBR ^TM^ Safe DNA gel stain (Thermo Fisher Scientific) for 40 minutes, at 200 V and 400 mA. The final gels were visualized with Azure Biosystems C600 gel imaging system Bio-Rad Laboratories, Inc.

### Assessment of *ompW and* multiplex PCR assay sensitivity

We assessed the sensitivity of our PCR assay with gBlocks at six target concentrations— 2,000, 1,000, 500, 250, 100, and 50 copies per reaction—using the optimized cycling time, annealing temperature, and primer concentration. Twelve (12) replicates were tested for each concentration prepared by performing serial dilution. To ensure that the DNA stock concentration matched the necessary copy number (for example, 1,000 copies/µL to achieve 2,000 copies in 2 µL), 2 µL of DNA template was added for each dilution level. Throughout the dilution processes, TE buffer was utilized as the diluent, and homogeneity was ensured by vigorous mixing with a vortex mixer. Thereafter, the PCR was run for each dilution, and the presence of visible bands on the gel indicated that the amplification was successful.

## Statistical Analysis

The optimization and validation of the PCR assay were qualitative. We utilized Quo data software specifically developed for the study of qualitative PCR experiments to assess the sensitivity of our assay. This program applies the probability of detection (POD) approach, as described by Uhlig and others (26), to estimate the 95% confidence interval (CI) for the limit of detection (LOD95%).The LOD95% is the average number of DNA copies needed in a sample to have a 95% chance of being detected. This is an important measure of how sensitive our test is at detecting the target. R programming statistical software version 4.5.0 was used to run a Two-sided Fisher’s exact test to check the statistical significance of all four assays (*ompW, O1-rfb, ctxA*, and *O139-rfb*) at each concentration level for both platforms. Holm–Bonferroni corrections were used on the p-values to account for the fact that each assay was tested more than once.

## Results

### Evaluation of new optimized PCR assay for *V. cholerae* Detection PCR Cycling time

Out of the three different cycling conditions tested for *ompW* assay, the best cycling condition was the first *ompW* cycling condition (*ompW*-1) with an initial denaturation step at 94°C for 5 min, followed by 30 cycles of denaturation (94°C, 30 sec), annealing (64°C, 30 sec), and extension (72°C, 30 sec), and a final extension step at 72°C for 7 min on both thermal platforms. Out of the four cycling conditions tested for the multiplex assay reaction, the baseline cycling condition (multiplex-1) with 1 min for denaturation, annealing and extension, worked best. Given that (*ompW*-1) and (multiplex-1) produced better and more equal bands on both thermal cyclers, they were both chosen for subsequent experiments.

### Annealing temperature

Different annealing temperatures were tested to find the highest possible temperature that produced the best equal bands for the *ompW* and multiplex (*ctxA*, 01-*rfb* and 0139-*rfb*) using both thermal cyclers. The annealing temperature on both thermocyclers’ platforms for *ompW* ranged from 63 °C to 68 °C while that of the multiplex assay ranged from 55°C to 60°C.

Bands for *ompW* were detected for the entire temperature range (63°C to 68°C) on both platforms. However, a difference was observed for the best visible bands for *ompW* on SimpliAmp was produced at 65°C, while for the Bio-Rad, the best visible bands were produced at 66°C. Given these results, 65°C and 66°C were chosen for subsequent *ompW* primer concentrations on SimpliAmp and Bio-Rad respectively.

Similar to the *ompW* assay, bands for all targets in our multiplex were detected across the 55–60 °C range. At high template concentrations (100,000 gBlock copies), the strongest bands were observed at 59 °C on the SimpliAmp and 60 °C on the Bio-Rad. However, at these temperatures bands were not detectable at the lower template concentration of 2,000 gBlock copies. In contrast, annealing at 55 °C and 56 °C allowed detection down to 2,000 copies; therefore, these temperatures were selected for subsequent primer concentration optimization.

### Uniplex *ompW* primer concentration

Different primer concentrations ranging from 1.0 µM to 0.6 µM were tested to determine the optimal concentration for the *ompW* assay on both platforms as shown in **Fig. 4**. Bands were visible at all concentrations, but at 0.9 µM the amplification was more consistent across duplicates and more uniform between the two thermal cyclers compared to 1.0 µM, where replicate variability was greater. Importantly, at 0.9 µM bands remained strong and detection was still possible down to 2,000 gBlock copies. For these reasons, 0.9 µM was selected for subsequent sensitivity testing.

**Fig. 4:**
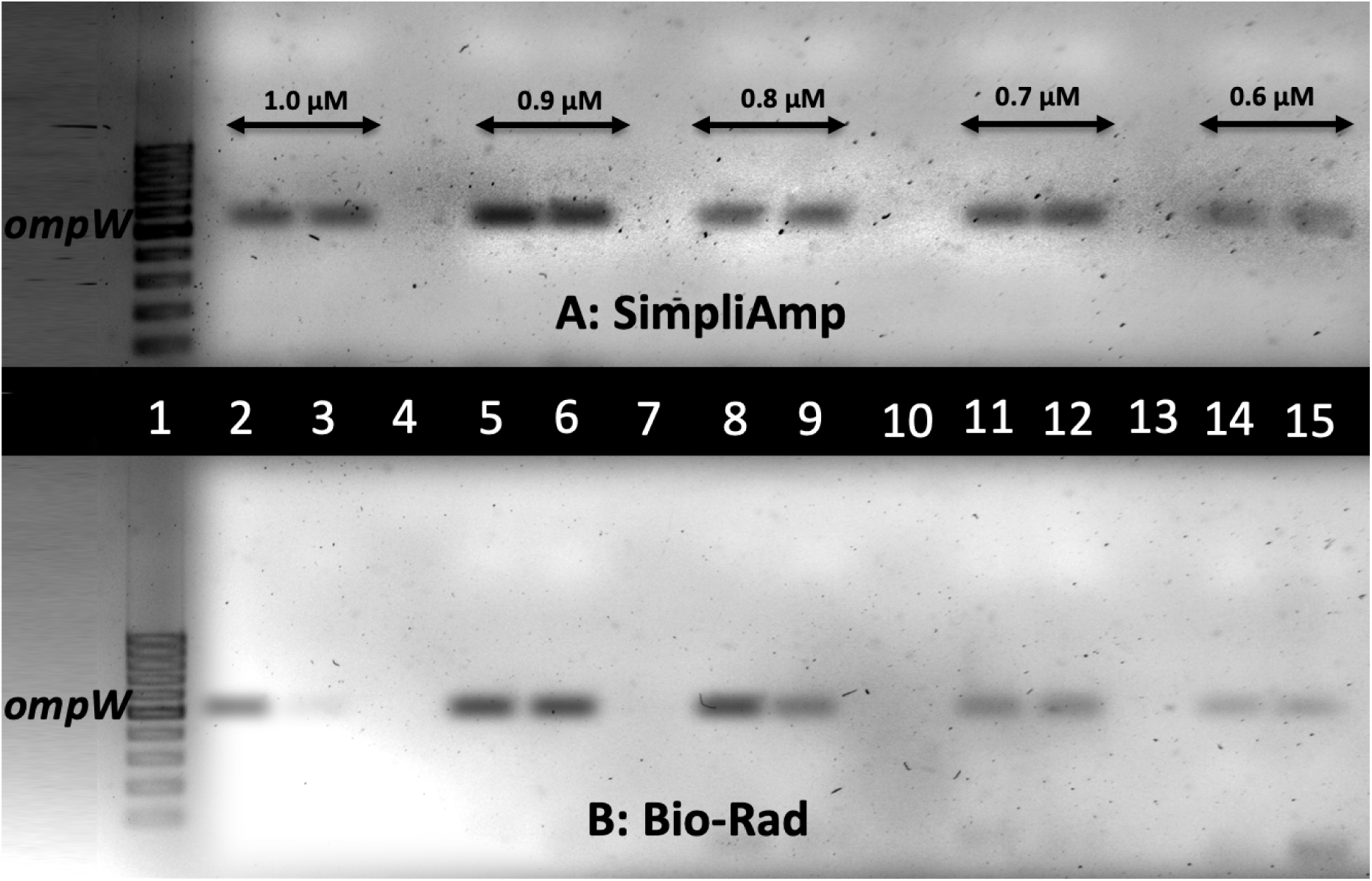
Gel electrophoresis of the ompW Uniplex PCR assay performed on the SimpliAmp (A) and Bio-Rad (B) platforms at different primer concentrations. Duplicate reactions were run for each concentration: 1.0 µM (lanes 2–3), 0.9 µM (lanes 5–6), 0.8 µM (lanes 8–9), 0.7 µM (lanes 11–12), and 0.6 µM (lanes 14–15).

### Multiplex primer concentration

For the multiplex assay, different primer concentration combinations were evaluated in a matrix format (**Table 4**). The optimal sensitivity was achieved using the 9th combination, where *ctxA* was set at 0.13 µM and both *O1-rfb* and *O139-rfb* at 1.00 µM. This combination was selected because it enabled the detection of all three biomarkers down to 250 copies, which was not achieved comparably with other tested combinations on both platforms (**Fig. 5 and 6)**. Notably, the baseline combination—*ctxA* at 0.17 µM and both *O1-rfb* and *O139-rfb* at 2.7 µM**—**failed to achieve the same sensitivity, at an annealing temperature of 55 °C (SimpliAmp,) and 56 °C (Bio-Rad). Summary of all optimized conditions are presented in **Table 5**.

**Fig. 5:**
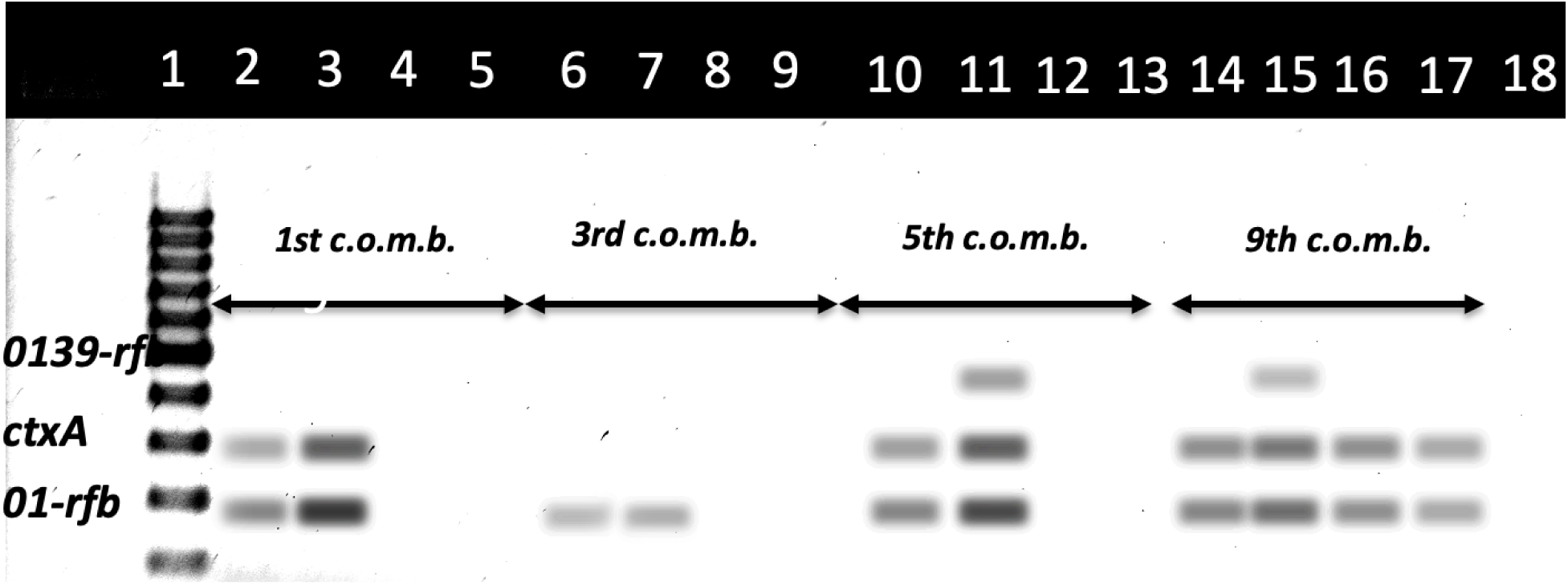
Gel electrophoresis of multiplex PCR products showing detection of O139-rfb, ctxA, and O1-rfb targets using the 1st, 3rd, 5th, and 9th primer combinations on the SimpliAmp (Applied Biosystems®). For each combination, duplicate reactions were performed with 250 gBlock copies (lanes 2–3, 6–7, 10–11, 14–15) and 100 gBlock copies (lanes 4–5, 8–9, 12–13, 16–17). Target bands are indicated on the left.

**Fig. 6:**
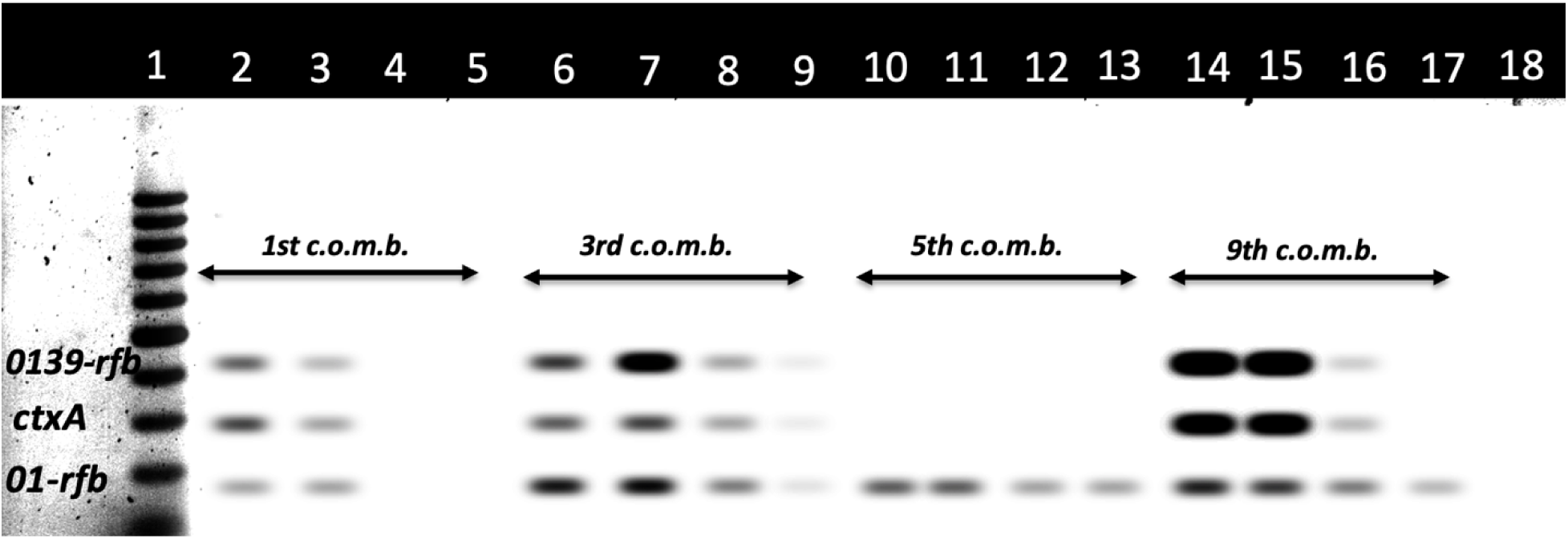
Gel electrophoresis of multiplex PCR products showing detection of O139-rfb, ctxA, and O1-rfb targets using the 1st, 3rd, 5th, and 9th primer combinations on the Bio-Rad (Bio-Rad Laboratories, Inc). For each combination, duplicate reactions were performed with 250 gBlock copies (lanes 2–3, 6–7, 10–11, 14–15) and 100 gBlock copies (lanes 4–5, 8–9, 12–13, 16–17). Target bands are indicated on the left.

**Table 5:**
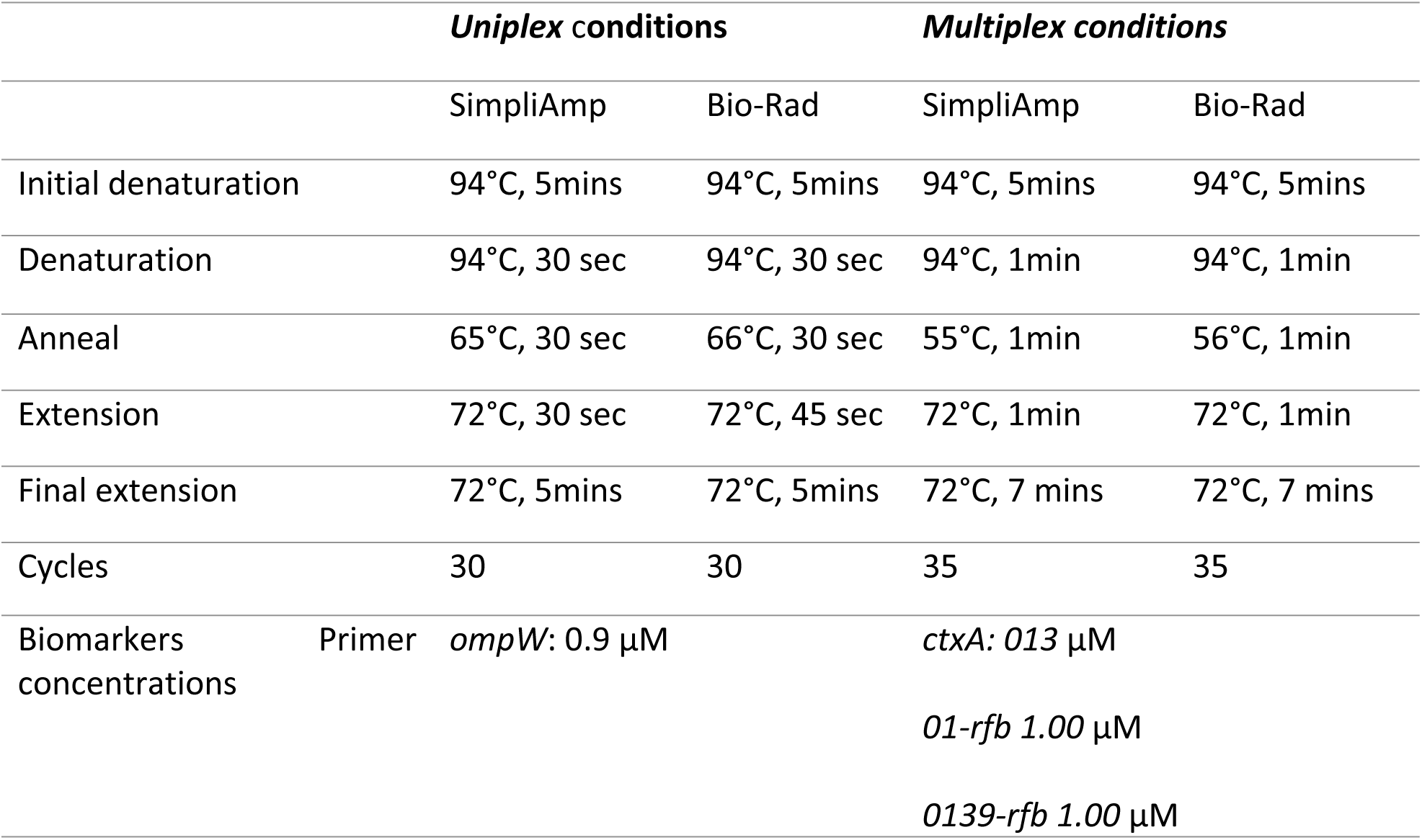
Summary of assays optimized conditions

## Summary of optimized conditions

### *ompW* assay sensitivity

The sensitivity of the *ompW* assay was evaluated using serial dilutions of 2,000, 1,000, 500, 250, 100, and 50 gBlock copies (bacteria), with twelve (12) replicates per dilution level. The *ompW* assay demonstrated identical performance on both the SimpliAmp by Applied Biosystem® and Bio-Rad platforms by Bio-Rad Laboratories, Inc. (**Table 6**). Both platforms achieved a 100% detection rate across 2000–250 copies. We observed a reduced detection at 100 copies (2/12 positives) while at 50 copies, no amplification was observed. Both platform LOD 95% = 523 copies with, 95% CI = [347.451, 785.784] (**Fig. 7**). There was no statistical difference (p=1.000) between platforms at any dilution level.

**Table 6:**
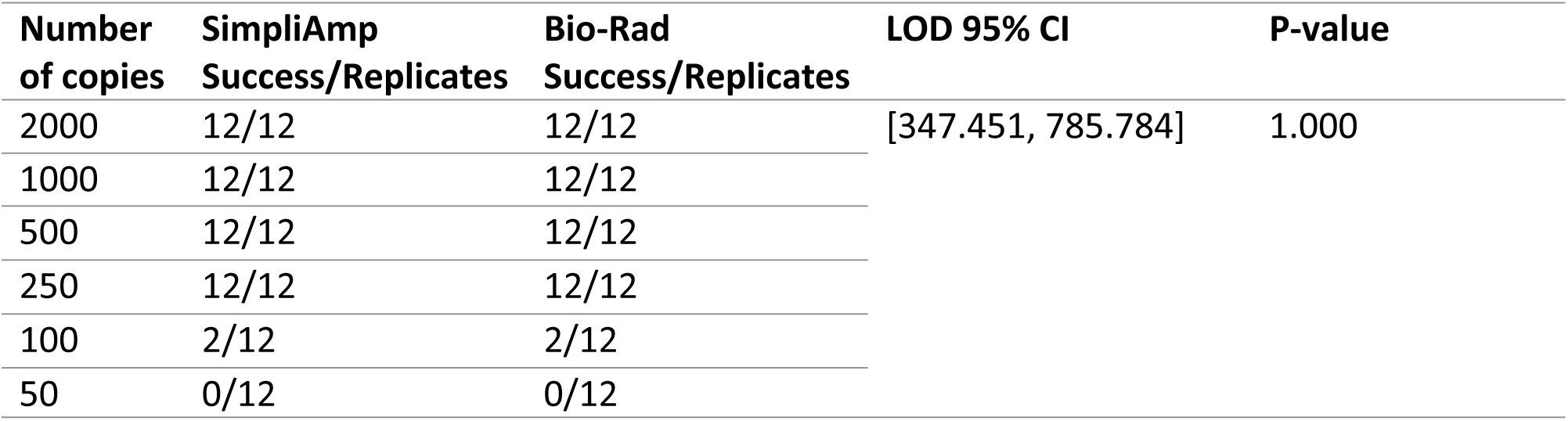
ompW sensitivity test using Both Thermal cyclers

**Fig. 7:**
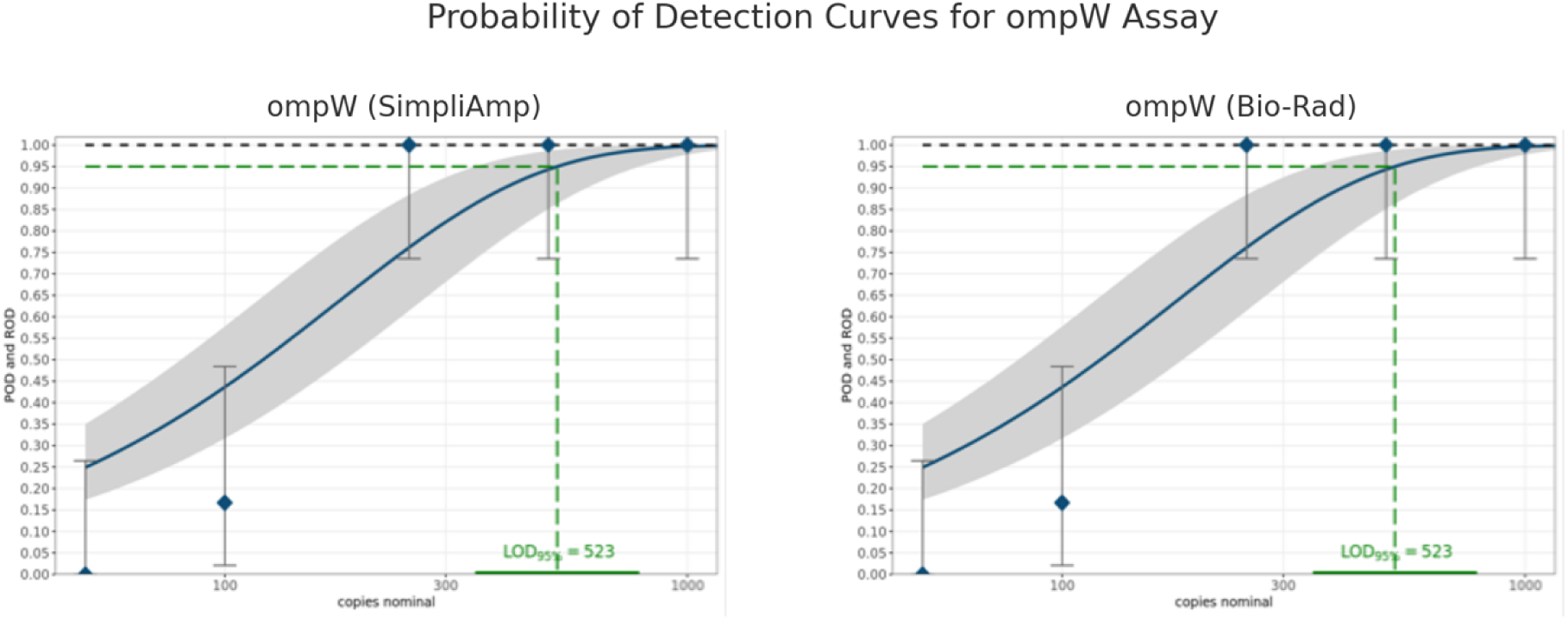
The figure shows the probability of detection (POD) curve generated from replicate testing of serially diluted ompW gBlock DNA (2,000–50 copies) for both platforms. Blue diamonds represent observed rates of detection (ROD) for each dilution level, while the blue line indicates the fitted mean POD curve with its 95% confidence range shaded in gray. The black dashed line represents the ideal POD curve (100% detection). The green dashed line denotes the estimated limit of detection at 95% confidence.

## Multiplex assay sensitivity

Similar to the *ompW* assay, the multiplex assay was evaluated using serial dilutions of 2,000, 1,000, 500, 250, 100, and 50 gBlock copies (bacteria), with twelve (12) replicates per dilution level. In **Table 7**, both platforms were able to detect all copies down to 1000 copies every time for the *O1-rfb* assay. Detection stayed high at 500 copies (11/12 on SimpliAmp vs. 12/12 on Bio-Rad). At 250 copies, the detection was a little better on SimpliAmp (10/12 vs. 9/12). SimpliAmp, on the other hand, demonstrated a trend toward improved sensitivity at lower concentrations, detecting 5/12 and 3/12 replicates at 100 and 50 copies, respectively, compared to Bio-Rad’s 2/12 and 0/12. Using the web-based software, SimpliAmp LOD 95% was estimated at 509 copies, with a 95% CI = [338.585, 765.733] (**Fig. 8**). For the Bio-Rad, the LOD95% of 333 copies, with 95% CI = [254.792, 687.973]. Despite the SimpliAmp showing greater sensitivity at 100 and 50 copies, the differences are not statistically significant at (p=0.371) and (p=0.217) respectively.

**Table 7:** 01-rfb sensitivity (LOD) test

| Number of copies | SimpliAmp Success/replicates | LOD 95% CI | Bio-Rad Success/Replicates | LOD 95% CI | P-Value |
| --- | --- | --- | --- | --- | --- |
| 2000 | 12/12 | [338.9, 765.7] | 12/12 | [254.8, 688.0] | 1.000 |
| 1000 | 12/12 |  | 12/12 |  |  |
| 500 | 11/12 |  | 12/12 |  |  |
| 250 | 10/12 |  | 9/12 |  |  |
| 100 | 5/12 |  | 2/12 |  | 0.371 |
| 50 | 3/12 |  | 0/12 |  | 0.217 |

**Fig. 8:**
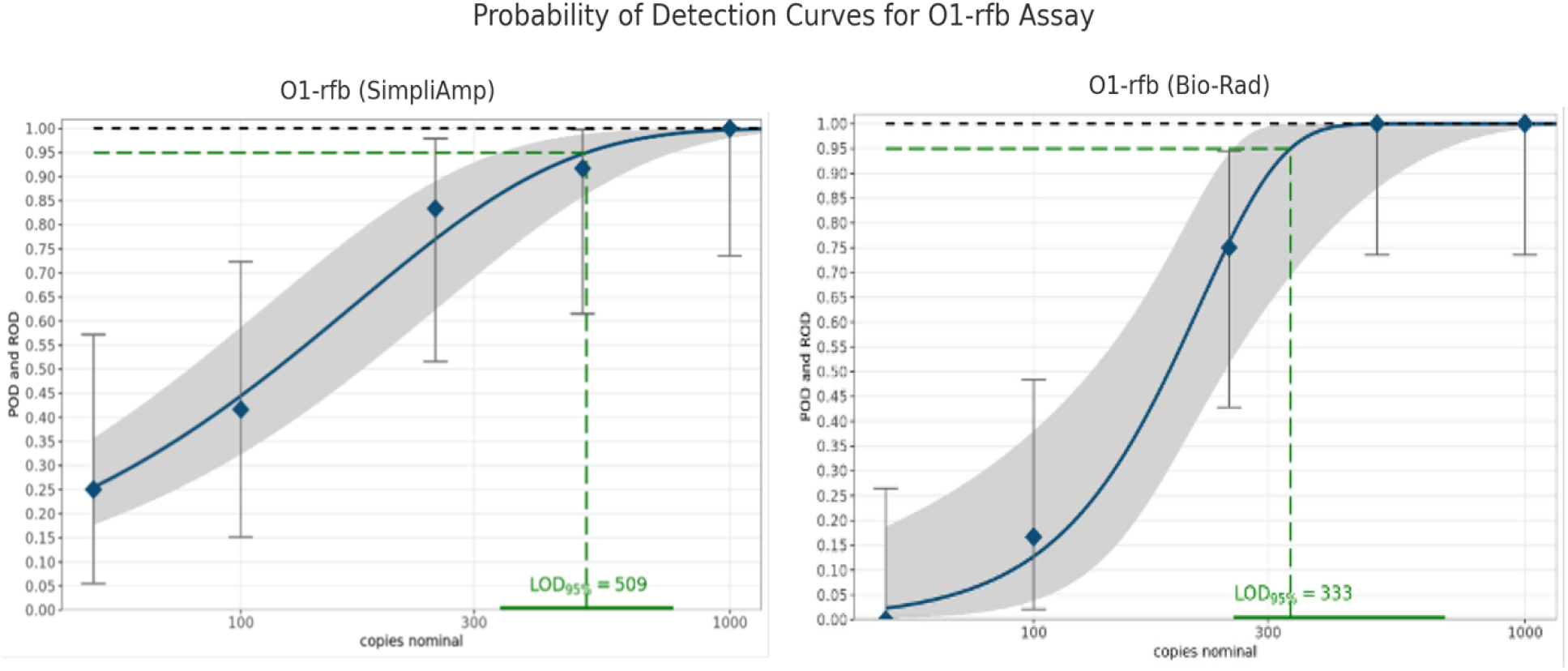
The figure shows the probability of detection (POD) curve generated from replicate testing of serially diluted 01-rfb gBlock DNA (2,000–50 copies) for both platforms. Blue diamonds represent observed rates of detection (ROD) for each dilution level, while the blue line indicates the fitted mean POD curve with its 95% confidence range shaded in gray. The black dashed line represents the ideal POD curve (100% detection). The green dashed line denotes the estimated limit of detection at 95% confidence.

For ctxA assay, both platforms detected all of the replicates (12/12) in 2000 and 1000 copies. However, at 500 copies, SimpliAmp platform was able to detect (10/12) compared to Bio-Rad which detected (5/12). Despite the higher successes on the SimpliAmp platform over Bio-Rad being at this concentration, the difference was not statistically significant (p = 0.089) as shown in **Table 8**. At 250 and 100 copies, detection was low on both platforms with successes at (3/12 vs. 4/12 and 2/12 vs. 1/12) on SimpliAmp and Bio-Rad respectively. At 50 copies, neither platform was amplified. SimpliAmp LOD95% was estimated at 684 copies (**Fig. 9**) with a 95% CI = [500.791, 1322.573]. The Bio-Rad LOD95% was estimated at 974 with 95% CI **=** [713.851, 1783.678].

**Table 8:** ctxA sensitivity test

| Number of copies | SimpliAmp Success/Replicates | LOD 95% CI | Bio-Rad Success/Replicates | LOD 95% CI | P-value |
| --- | --- | --- | --- | --- | --- |
| 2000 | 12/12 | [500.8, 1322.6] | 12/12 | [713.9, 1783.7] | 1.000 |
| 1000 | 12/12 |  | 12/12 |  |  |
| 500 | 10/12 |  | 5/12 |  |  |
| 250 | 3/12 |  | 4/12 |  |  |
| 100 | 2/12 |  | 1/12 |  |  |
| 50 | 0/12 |  | 0/12 |  |  |

**Fig. 9:**
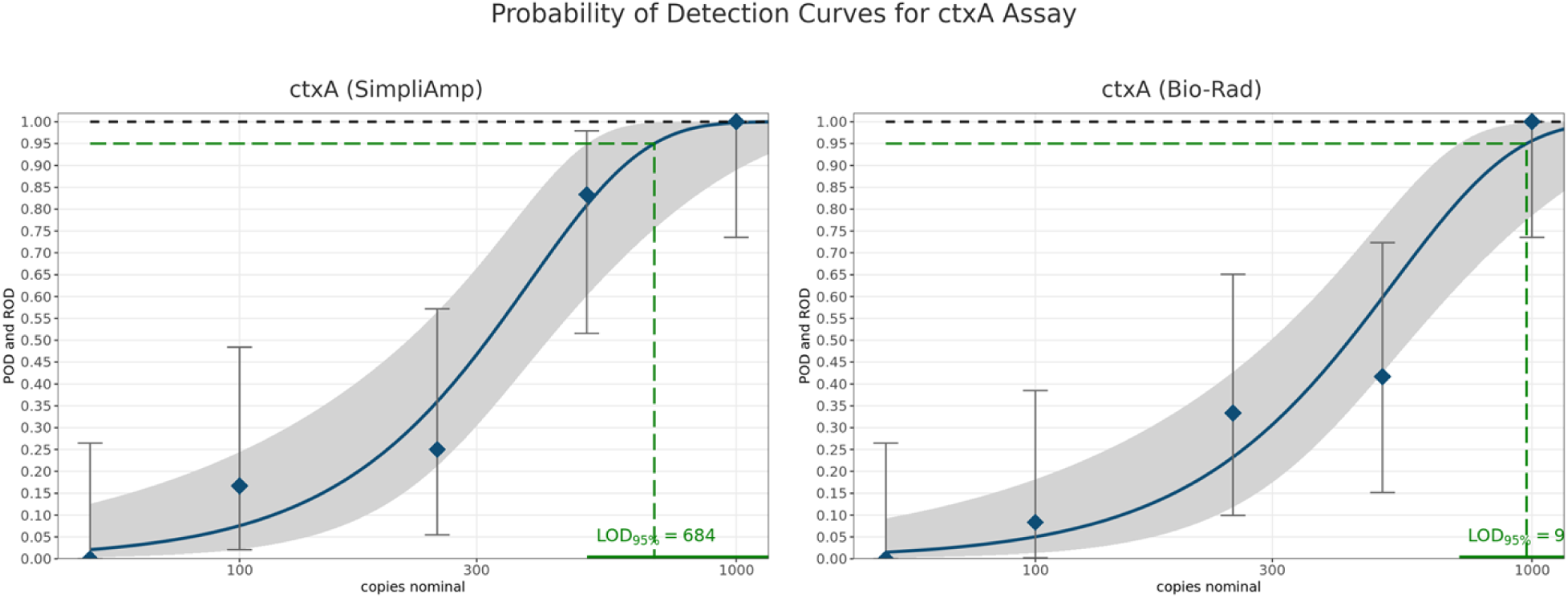
The figure shows the probability of detection (POD) curve generated from replicate testing of serially diluted ctxA gBlock DNA (2,000–50 copies) for both platforms. Blue diamond’s represent observed rates of detection (ROD) for each dilution level, while the blue line indicates the fitted mean POD curve with its 95% confidence range shaded in gray. The black dashed line represents the ideal POD curve (100% detection). The green dashed line denotes the estimated limit of detection at 95% confidence.

Similar to the other assays, both platforms detected all the replicates (12/12) in 2000 and 1000 copies for the O139-rfb assay as shown in **Table 9**. At 500 copies, similar detection (10/12 vs. 8/12) was observed on SimpliAmp and Bio-Rad respectively. Bio-Rad detected more copies at lower concentrations, detecting 7 out of 12 and 3 out of 12 at 250 and 100 copies, while SimpliAmp only found 2 out of 12 and 1 out of 12. Even while these differences weren’t statistically significant, they do show a tendency toward Bio-Rad being more sensitive with fewer template numbers. SimpliAmp LOD95% was estimated at 639 copies (**Fig. 10)** with a 95% CI = [489.817, 1251.309]. The Bio-Rad LOD95% was estimated at 1060 with 95% CI = [719.143, 1561.653].

**Table 9:** 0139-rfb sensitivity test

| Number of copies | SimpliAmp Success/Replicates | LOD 95% CI | Bio-Rad Success/Replicates | LOD 95% CI | P-value |
| --- | --- | --- | --- | --- | --- |
| 2000 | 12/12 | [489.8,1251.3] | 12/12 | [719.1,1561.7] | 1.000 |
| 1000 | 12/12 |  | 12/12 |  |  |
| 500 | 10/12 |  | 8/12 |  | 0.640 |
| 250 | 2/12 |  | 7/12 |  | 0.089 |
| 100 | 1/12 |  | 3/12 |  | 0.590 |
| 50 | 0/12 |  | 0/12 |  | 1.000 |

**Fig. 10:**
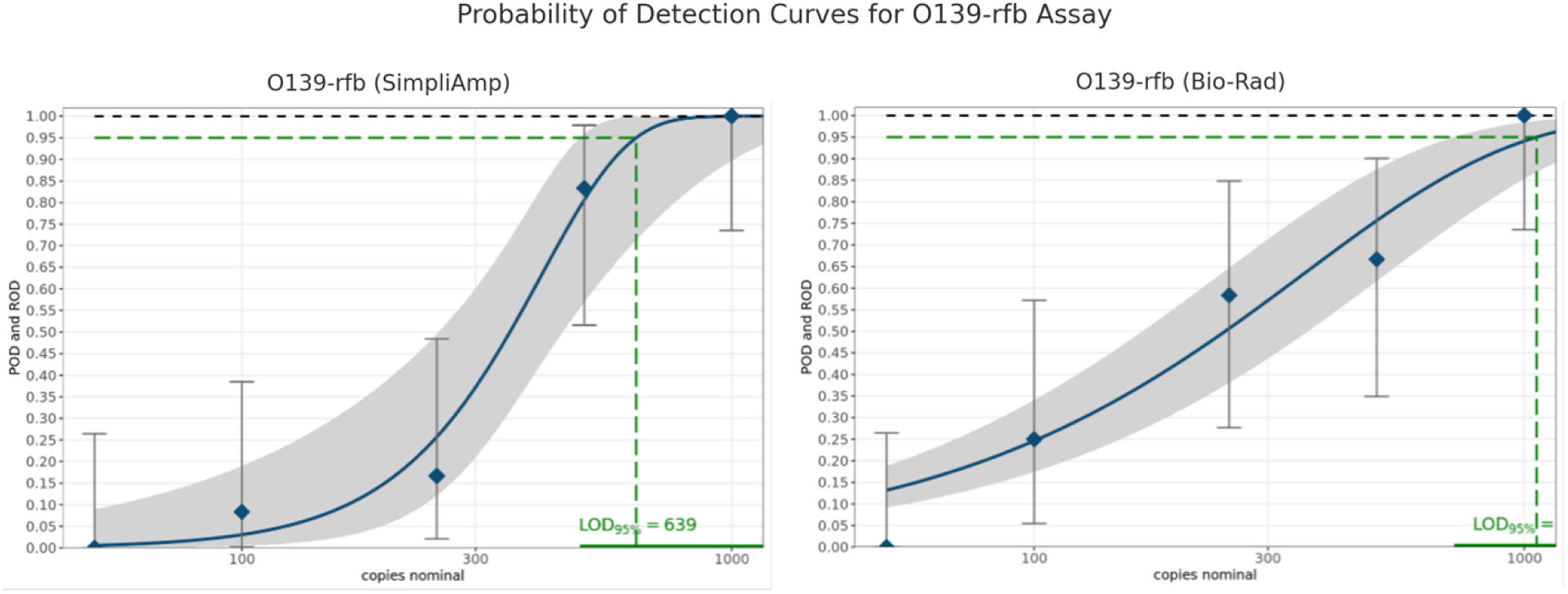
The figure shows the probability of detection (POD) curve generated from replicate testing of serially diluted 0139-rfb gBlock DNA (2,000–50 copies) for both platforms. Blue diamonds represent observed rates of detection (ROD) for each dilution level, while the blue line indicates the fitted mean POD curve with its 95% confidence range shaded in gray. The black dashed line represents the ideal POD curve (100% detection). The green dashed line denotes the estimated limit of detection at 95% confidence.

Similar to the other assays, both platforms detected all the replicates (12/12) in 2000 and 1000 copies for the O139-rfb assay as shown in **Table 9**. At 500 copies, similar detection (10/12 vs. 8/12) was observed on SimpliAmp and Bio-Rad respectively. Bio-Rad detected more copies at lower concentrations, detecting 7 out of 12 and 3 out of 12 at 250 and 100 copies, while SimpliAmp only found 2 out of 12 and 1 out of 12. Even while these differences weren’t statistically significant, they do show a tendency toward Bio-Rad being more sensitive with fewer template numbers. SimpliAmp LOD95% was estimated at 639 copies (**Fig. 10)** with a 95% CI = [489.817, 1251.309]. The Bio-Rad LOD95% was estimated at 1060 with 95% CI = [719.143, 1561.653].

## Discussion

The availability of rapid diagnostic technologies for infectious diseases that are cheaper for Low-and Middle-Income Countries (LMICs) will mark a major step in improving global public health (27). The major challenge to this effort, however, is the majority of the population that needs these technologies resides in places with limited lab infrastructures (28–30). In this study, we compared effectiveness and sensitivity of two platforms-the Bio-Rad C1000 Touch which is priced at $12,160.00 (31) versus the SimpliAmp priced at $6,500.00 (32) to inform decision making in diagnostics since more expensive equipment are sometimes thought to be more efficient. To achieve this, different cycling time, annealing temperatures, and primer concentrations were tested for optimization on both platforms, and subsequent assay sensitivity was determined. For *ompW* assay, annealing temperature was optimized from 64°C in the baseline (23), to 65°C and 66°C on SimpliAmp and Bio-Rad respectively. Despite not being able to optimize annealing temperature for our multiplex assay from the baseline of 55°C (25) on the SimpliAmp platform, that was not the case on the Bio-Rad as our annealing temperature was optimized to 56°C.

Given that previous studies have shown that an increase in annealing temperature improved specificity(33,34), any increase in annealing temperature, as applied in this study, improved the specificity of the assay. This was particularly valuable for LMICs, where false-positive results could lead to unnecessary exposure to disease and further use of limited research supplies (35). Our study also shows that sensitivity is maintained within a defined temperature range and not one specific temperature. In LMICs, where laboratory conditions might not always permit very exact thermal control, this adaptability is particularly vital. By demonstrating that efficient detection of *V. cholerae* is still possible over a range of annealing temperatures, this study supports the possibility of robust, field-adaptable PCR protocols that can withstand slight variations in conditions.

Despite both thermal cycler platforms detecting *ompW* at 100,000 gBlocks copies, only higher primer concentrations of 1.00 µM-0.8 µM, could detect *ompW* down to 2000 gBlocks copies or lower. The original study for baseline conditions did not report its assay LOD values, it only used the PCR assay to find *V. cholerae* strains (23). However, another study that developed a single multiplex PCR assay to detect ompW in environmental samples reported a sensitivity of 5 × 10⁴ bacterial cells, meaning that at least 50,000 *V. cholerae* cells had to be present in a sample for the ompW gene to be detected (36). However, in our study, the *ompW* assay was sensitive enough to detect *V. cholerae* at 100 gBlocks copies or bacterial cells on both thermal cycler platforms. This might be due to the pure concentration of target gBlocks used in our study versus the enrichment step used in environmental samples.

Using our selected combination for the multiplex assay, SimpliAmp was sensitive enough to detect *01-rfb* at 50 copies of gBlocks compared to 100 copies of *01-rfb* gBlocks on the Bio-Rad platform. Both thermal cyclers platforms detected *ctxA* and *0139-rfb* at 100 gBlocks copies. Our results are comparable with the study that was used to establish baseline conditions (24), which developed and evaluated a multiplex PCR assay for rapid detection of toxigenic *V. cholerae* where O1 and O139 were detected at 65 cfu and 200 cfu respectively (37). Our study demonstrates that the optimized assays perform reliably on both platforms, with SimpliAmp showing comparable or slightly better sensitivity than the Bio-Rad despite its lower cost. This makes the SimpliAmp thermal cycler a reliable and more affordable alternative to the Bio-Rad in LMICs and low-resources settings where cholera is endemic to support outbreak investigation and surveillance.

## Limitation of Study

One drawback of this study is that the PCR assay was evaluated in a controlled laboratory setting, which may not fully represent the intricacies of real-world applications. This method allowed for precise adjustments and assessments of assay performance, but it may not be the same in real-world scenarios, as environmental samples exhibit greater variability compared to laboratory samples. So, even though the bench study gives strong evidence for how well the assay works, more testing with different samples that have lots of inhibitors is needed to determine its detection limits and sensitivity in environmental samples.

## Conclusion

While cycling times from the baseline studies for both assays were maintained in the current study, annealing temperature was improved for the ompW assay on both thermal cycler platforms and for the multiplex assay on the Bio-Rad. This study also reduced primer concentrations for both assays from 1.0 µM to 0.9 µM and still achieved comparable results to other studies. Although this reduction was small, it represents a 10% decrease in primer usage. Over hundreds of reactions, this translates to significant savings in reagent costs, which is particularly important in resource-limited laboratories. The cost-effective SimpliAmp showed promise, as its performance was comparable to that of the Bio-Rad C1000. In LMICs, where access to standard PCR is often restricted by financial and infrastructural constraints, these refinements build upon existing knowledge and offer a workable solution for improving disease surveillance on reasonably cost-effective thermocyclers.

This study promotes the development of locally flexible and scalable diagnostic methods, which are essential for enhancing epidemic readiness and fostering resilience in LMIC public health systems, particularly in areas where cholera is endemic. Finally, this work adds to efforts to fill diagnostic gaps and guarantee that lifesaving, cost-effective instruments are available for use, especially for the most vulnerable communities.

## Author Contributions

Conceptualization, Aminata Kilungo, Gerardo Lopez, Khai Truong, Victor Okpanachi, Methodology, Gerardo Lopez, Khai Truong, Kerry Cooper, Aminata Kilungo Funding acquisition, Aminata Kilungo; Project administration, Aminata Kilungo; Resources, Aminata Kilungo; Supervision, Aminata Kilungo, Gerardo Lopez; Writing – original draft, Victor Okpanachi and Aminata Kilungo; Visualization, Victor Okpanachi; Writing – review & editing, Victor Okpanachi, Aminata Kilungo, Gerardo Lopez, Khai Truong, and Kerry Cooper

## References

1. Kanungo S, Azman AS, Ramamurthy T, Deen J, Dutta S. Cholera. The Lancet. 2022 Apr;399(10333):1429–40.

2. Montero DA, Vidal RM, Velasco J, George S, Lucero Y, Gómez LA, et al. Vibrio cholerae, classification, pathogenesis, immune response, and trends in vaccine development. Front Med. 2023 May 5;10:1155751.

3. Mehrabadi JF, Morsali P, Nejad HR, Imani Fooladi AA. Detection of toxigenic Vibrio cholerae with new multiplex PCR. Journal of Infection and Public Health. 2012 June;5(3):263–7.

4. Moore S, Thomson N, Mutreja A, Piarroux R. Widespread epidemic cholera caused by a restricted subset of Vibrio cholerae clones. Clinical Microbiology and Infection. 2014 May;20(5):373–9.

5. Asantewaa AA, Odoom A, Owusu-Okyere G, Donkor ES. Cholera Outbreaks in Low- and Middle-Income Countries in the Last Decade: A Systematic Review and Meta-Analysis. Microorganisms. 2024 Dec 4;12(12):2504.

6. Yan Y, Zhan L, Zhu G, Zhang J, Li P, Chen L, et al. Direct and Rapid Identification of Vibrio Cholerae Serogroup and Toxigenicity by a Novel Multiplex Real-Time Assay. Pathogens. 2022 July 30;11(8):865.

7. Xiao L, Li Z, Dou X, Long Y, Li Z, Liu Y, et al. Advancing microbial risk assessment: perspectives from the evolution of detection technologies. NPJ Sci Food. 2025 July 28;9(1):157.

8. Morshed MG, Lee MK, Jorgensen D, Isaac-Renton JL. Molecular methods used in clinical laboratory: prospects and pitfalls. FEMS Immunology & Medical Microbiology. 2007 Mar;49(2):184–91.

9. Park SH, Hwang KA, Ahn JH, Nam JH. Evaluation of Multiplex Polymerase Chain Reaction Assay for the Simultaneous Detection of Sexually Transmitted Infections Using Swab Specimen. J Bacteriol Virol. 2020;50(1):44.

10. Department of Pediatrics, Faculty of Medicine, Can Tho University of Medicine and Pharmacy, Can Tho City, Vietnam, Tran KQ, Pham VH, Laboratory of Nam Khoa Biotek Company, Vietnam Research and Development Institute of Clinical Microbiology, Ho Chi Minh City, Vietnam, Vo CM, Department of Pediatrics, Faculty of Medicine, Can Tho University of Medicine and Pharmacy, Can Tho City, Vietnam, et al. Comparison of Real-time Polymerase Chain Reaction and Culture for Targeting Pathogens in Pediatric Severe Community-Acquired Pneumonia. Turkish Archives of Pediatrics. 2024 Aug 2;59(4):383–9.

11. CDC. Escherichia coli, Diarrheagenic | CDC Yellow Book 2024 [Internet]. [cited 2025 Apr 18]. Available from: https://www.nc.cdc.gov/travel/yellowbook/2024/infections-diseases/escherichia-coli-diarrheagenic

12. Hounmanou YMG, Leekitcharoenphon P, Hendriksen RS, Dougnon TV, Mdegela RH, Olsen JE, et al. Surveillance and Genomics of Toxigenic Vibrio cholerae O1 From Fish, Phytoplankton and Water in Lake Victoria, Tanzania. Front Microbiol. 2019 Apr 30;10:901.

13. Naha A, Chowdhury G, Ghosh-Banerjee J, Senoh M, Takahashi T, Ley B, et al. Molecular Characterization of High-Level-Cholera-Toxin-Producing El Tor Variant Vibrio cholerae Strains in the Zanzibar Archipelago of Tanzania. J Clin Microbiol. 2013 Mar;51(3):1040–5.

14. Schoonbroodt S, Ichanté JL, Boffé S, Devos N, Devaster JM, Taddei L, et al. Real-time PCR has advantages over culture-based methods in identifying major airway bacterial pathogens in chronic obstructive pulmonary disease: Results from three clinical studies in Europe and North America. Front Microbiol. 2023 Feb 24;13:1098133.

15. O’Leary J, Corcoran D, Lucey B. Comparison of the EntericBio Multiplex PCR System with Routine Culture for Detection of Bacterial Enteric Pathogens. J Clin Microbiol. 2009 Nov;47(11):3449–53.

16. Courtney JW, Kostelnik LM, Zeidner NS, Massung RF. Multiplex Real-Time PCR for Detection of *Anaplasma phagocytophilum* and *Borrelia burgdorferi*. J Clin Microbiol. 2004 July;42(7):3164–8.

17. Gubala AJ, Proll DF. Molecular-beacon multiplex real-time PCR assay for detection of Vibrio cholerae. Appl Environ Microbiol. 2006 Sept;72(9):6424–8.

18. Waturangi DE, Amadeus S, Kelvianto YE. Survival of enteroaggregative Escherichia coli and Vibrio cholerae in frozen and chilled foods. J Infect Dev Ctries. 2015 Aug 29;9(8):837–43.

19. Marquet F, Mora N, Incani RN, Jesus J, Méndez N, Mujica R, et al. Comparison of different PCR amplification targets for molecular diagnosis of *Strongyloides stercoralis*. J Helminthol. 2023;97:e88.

20. Al-Sa’ady AT, Baqer KA, Al-Salim ZKS. Molecular detection and phylogenetic analysis of Vibrio cholerae genotypes in Hillah, Iraq. New Microbes and New Infections. 2020 Sept;37:100739.

21. Ontweka LN, Deng LO, Rauzier J, Debes AK, Tadesse F, Parker LA, et al. Cholera Rapid Test with Enrichment Step Has Diagnostic Performance Equivalent to Culture. PLoS One. 2016;11(12):e0168257.

22. Chen YL, Lin CC, Yang SC, Chen WL, Chen JR, Hou YH, et al. Five Technologies for Detecting the EGFR T790M Mutation in the Circulating Cell-Free DNA of Patients With Non-small Cell Lung Cancer: A Comparison. Front Oncol. 2019 July 16;9:631.

23. Nandi B, Nandy RK, Mukhopadhyay S, Nair GB, Shimada T, Ghose AC. Rapid Method for Species-Specific Identification of *Vibrio cholerae* Using Primers Targeted to the Gene of Outer Membrane Protein OmpW. J Clin Microbiol. 2000 Nov;38(11):4145–51.

24. Alam M, Sultana M, Nair GB, Sack RB, Sack DA, Siddique AK, et al. Toxigenic *Vibrio cholerae* in the Aquatic Environment of Mathbaria, Bangladesh. Appl Environ Microbiol. 2006 Apr;72(4):2849–55.

25. Dalusi L, Lyimo TJ, Lugomela C, Hosea KMM, Sjöling S. Toxigenic Vibrio cholerae identified in estuaries of Tanzania using PCR techniques. FEMS Microbiology Letters [Internet]. 2015 Mar 1 [cited 2025 Jan 18];362(5). Available from: https://academic.oup.com/femsle/article/doi/10.1093/femsle/fnv009/468448

26. Uhlig S, Frost K, Colson B, Simon K, Mäde D, Reiting R, et al. Validation of qualitative PCR methods on the basis of mathematical–statistical modelling of the probability of detection. Accred Qual Assur. 2015 Apr;20(2):75–83.

27. Wong G, Wong I, Chan K, Hsieh Y, Wong S. A Rapid and Low-Cost PCR Thermal Cycler for Low Resource Settings. PLoS One. 2015;10(7):e0131701.

28. Chin CD, Linder V, Sia SK. Lab-on-a-chip devices for global health: Past studies and future opportunities. Lab Chip. 2007;7(1):41–57.

29. Koudokpon H, Lègba B, Sintondji K, Kissira I, Kounou A, Guindo I, et al. Empowering public health: building advanced molecular surveillance in resource-limited settings through collaboration and capacity-building. Front Health Serv. 2024 June 18;4:1289394.

30. Aborode AT, Adesola RO, Scott GY, Arthur-Hayford E, Otorkpa OJ, Kwaku SD, et al. Bringing lab to the field: Exploring innovations in point-of-care diagnostics for the rapid detection and management of tropical diseases in resource-limited settings. Advances in Biomarker Sciences and Technology. 2025;7:28–43.

31. Choose Your Cycler – Save Big on Bio-Rad Thermal Cyclers | Bio-Rad [Internet]. [cited 2025 Sept 25]. Available from: https://www.bio-rad.com/en-us/life-science-research/promotions/choose-your-cycler-save-big-bio-rad-thermal-cyclers

32. Applied Biosystems SimpliAmp Thermal Cycler - PCR Equipment and Supplies, Thermal Cyclers [Internet]. [cited 2025 Sept 25]. Available from: https://www.fishersci.com/shop/products/simpliamp-thermal-cycler-extended-warranty-package-5/A24811

33. Malhotra K. Interaction and effect of annealing temperature on primers used in differential display RT-PCR. Nucleic Acids Research. 1998 Feb 1;26(3):854–6.

34. Latham S, Hughes E, Budgen B, Morley A. Inhibition of the PCR by genomic DNA. Kalendar R, editor. PLoS ONE. 2023 Apr 21;18(4):e0284538.

35. Healy B, Khan A, Metezai H, Blyth I, Asad H. The impact of false positive COVID-19 results in an area of low prevalence. Clin Med (Lond). 2021 Jan;21(1):e54–6.

36. Goel AK, Ponmariappan S, Kamboj DV, Singh L. Single multiplex polymerase chain reaction for environmental surveillance of toxigenic—Pathogenic O1 and Non-O1vibrio cholerae. Folia Microbiol. 2007 Jan;52(1):81–5.

37. Hoshino K, Yamasaki S, Mukhopadhyay AK, Chakraborty S, Basu A, Bhattacharya SK, et al. Development and evaluation of a multiplex PCR assay for rapid detection of toxigenic *Vibrio cholerae* O1 and O139. FEMS Immunol Med Microbiol. 1998 Mar;20(3):201–7.

